# Motile ghosts of the halophilic archaeon, *Haloferax volcanii*

**DOI:** 10.1101/2020.01.08.899351

**Authors:** Yoshiaki Kinosita, Nagisa Mikami, Zhengqun Li, Frank Braun, Tessa EF. Quax, Chris van der Does, Robert Ishmukhametov, Sonja-Verena Albers, Richard M. Berry

## Abstract

Motility is seen across all domains of life^1^. Prokaryotes exhibit various types of motilities, such as gliding, swimming, and twitching, driven by supramolecular motility machinery composed of multiple different proteins^2^. In archaea only swimming motility is reported, driven by the archaellum (archaeal flagellum), a reversible rotary motor consisting of a torque-generating motor and a helical filament which acts as a propeller^3,4^. Unlike the bacterial flagellar motor (BFM), adenosine triphosphate (ATP) hydrolysis probably drives both motor rotation and filamentous assembly in the archaellum^5,6^. However, direct evidence is still lacking due to the lack of a versatile model system. Here we present a membrane-permeabilized ghost system that enables the manipulation of intracellular contents, analogous to the triton model in eukaryotic flagella^7^ and gliding *Mycoplasma*^8,9^. We observed high nucleotide selectivity for ATP driving motor rotation, negative cooperativity in ATP hydrolysis and the energetic requirement for at least 12 ATP molecules to be hydrolyzed per revolution of the motor. The response regulator CheY increased motor switching from counterclockwise (CCW) to clockwise (CW) rotation, which is the opposite of a previous report^10^. Finally, we constructed the torque-speed curve at various [ATP]s and discuss rotary models in which the archaellum has characteristics of both the BFM and F_1_-ATPase. Because archaea share similar cell division and chemotaxis machinery with other domains of life^11,12^, our ghost model will be an important tool for the exploration of the universality, diversity, and evolution of biomolecular machinery.

---

The archaellar motor has no homology with the BFM, but is evolutionarily and structurally related to bacterial type IV pili (T4P) for surface motility^3^. In Euryarchaeota, the filament is encoded by two genes, *flgA (flaA in Methanococcus)* and *flgB (flaB in Methanococcus)*, and the motor eight *flaC-J* (see Ref. 3 for details in Crenarchaeota). Euryarchaeota encode the full set of a chemotaxis system, *cheA, B, C, D, R, W,* and *Y*, like flagellated bacteria, which might have been acquired by horizontal gene transfer from *Bacillus/Thermotoga* groups^11^.

Figure 1a (top) shows the current association of functions with the motor genes, based on analysis of mutants and biochemical data: FlaC/D/E as switching proteins for the directional switch of archaellum rotation coupled with the signals from the chemotaxis system^13^; FlaG and FlaF complex interacting with the surface layer (S-layer), with FlaF regulating FlaG filament assembly^14,15^; FlaH as a regulator of the switch between assembly of the archaella and rotation^16^; FlaI as the ATP-driven motor for both assembly and rotation^5^; FlaJ as the membrane-spanning component. An inhibitor of proton translocating ATP synthases reduced both intracellular [ATP] and swimming speed in *Halobacterium salinarum*^6^, suggesting that archaellar rotation is driven by ATP hydrolysis at FlaI. However, direct evidence is lacking due to the lack of a reconstituted system. There is also no direct evidence as to which components are anchored to the cell and which rotate with the filament: Figure 1b illustrates possibilities which we discuss below.

**Figure 1.**
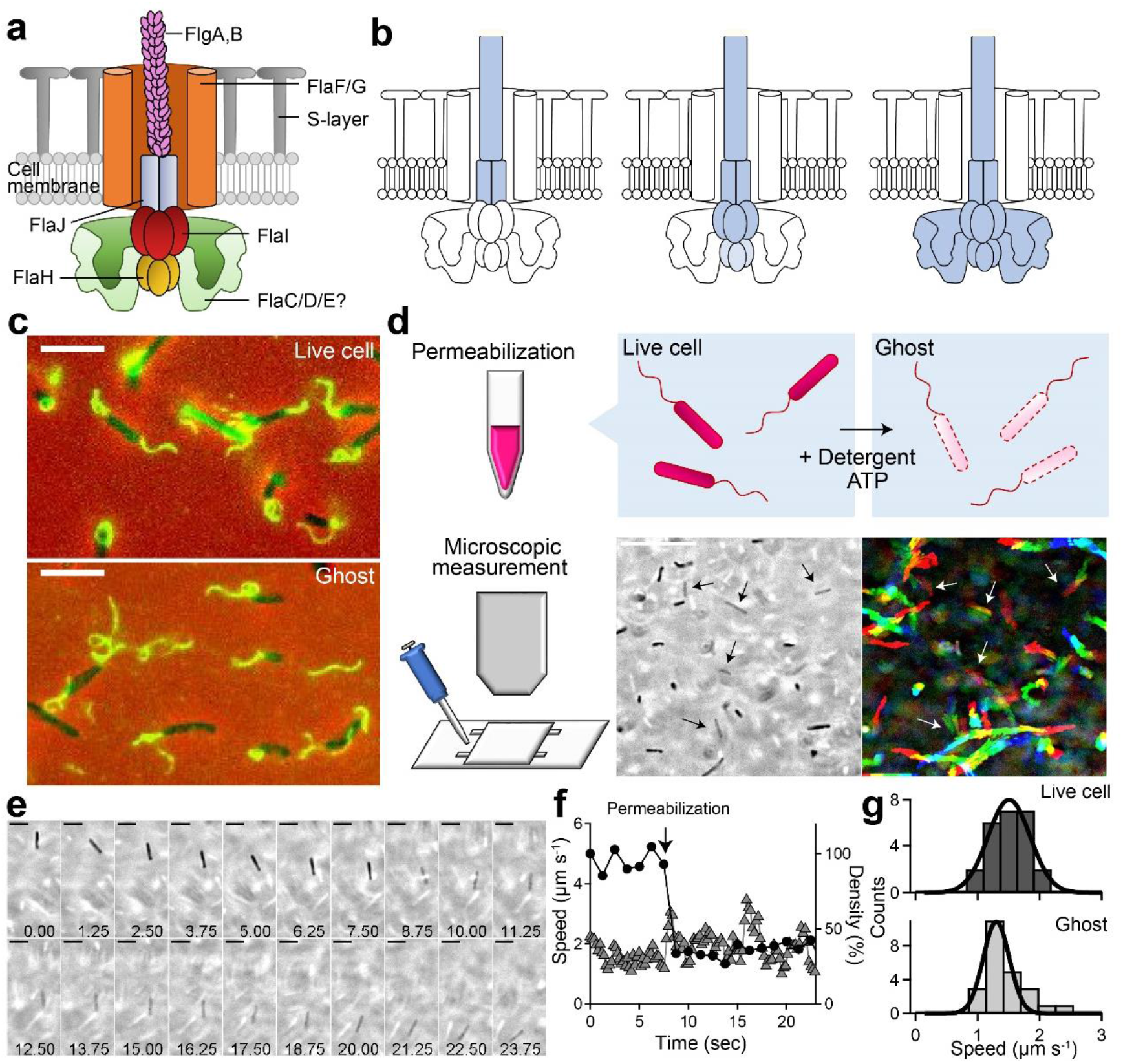
Swimming ghosts. (a) The current model of the archaeallar motor in Euryarchaeota (details in main text). (b) Different possibilities for which components of the archaellum are fixed relative to the cell (stator, white) and which rotate with the filament (rotor, blue). Details are described in the main text. (c) Merged phase-contrast and fluorescent images of live cells *(top)* and ghosts *(bottom)* labeled with streptavidin-dylight488 which binds to biotinylated filaments. (d) Procedures to observe swimming ghosts. Live cells were permeabilized in a tube, then induced into a flow chamber to observe swimming motility. *Lower middle:* Phase-contrast image. Black arrows indicate ghosts. Scale bar, 10 μm. Lower right: Sequential images with 0.5-s intervals, integrated for 30 sec with the intermittent color code “red → yellow → green → cyan → blue.” White arrows indicate trajectories of ghosts. (e) Sequential images of a change from a live cell to ghost. Scale bar, 5 μm. (f) Time course of swimming speed (triangles) and cell optical density (circles) from (d). Arrow indicates the time of permeabilization. (g) Histograms of swimming speed, 1.51 ± 0.34 μm s^-1^ before permeabilization (live cells) and 1.30 ± 0.21 μm s^-1^ after permeabilization (ghosts), (mean ± SD, n = 24).

Here we present an *in vitro* experimental system for the archaellum, similar to the Triton model for the eukaryotic flagellum^7^ and the permeabilized ghost model for gliding *Mycoplasma mobile*^8,9^ We use the halophilic archaeon *Haloferax volcanii. Hfx. volcanii* possesses multiple polar archaella and swims at 2-4 μm s^-1^ at room temperature, with CW rotation more efficient for propulsion than CCW (Fig. 1c *top,* Supplementary Result 1 and Supplementary Video 1)^17^ We increased the fraction of swimming *Hfx. volcanii* cells from 20-30 % to 80 % by adding 20 mM CaCl_2_ (Supplementary Figure 1a).

To prepare our experimental model system, we suspended motile cells in buffers containing detergent (0.015 % sodium cholate) and 2.5 mM ATP (Fig. 1d). Fluorescent imaging revealed that ghosts still possessed archaellar filaments, the cell membrane, and S-layer (Fig. 1c *bottom* and Supplementary Figure 2). The detergent reduced the refractive index of cells, indicating permeabilization of the cell membrane and corresponding loss of cytoplasm. Remarkably, the permeabilized cells still swam (Supplementary Video 2 and Fig. 1d *lower right)* We named them “ghosts,” as in similar experiments on *Mycoplasma mobile*^9^. Fig. 1e shows a typical example of a live swimming cell changing to a ghost, marked by a sudden change of image density at 8.75 sec. The solution contained 2.5 mM ATP and the swimming speed did not change dramatically when this cell became a ghost (Fig. 1f, see Supplementary Figure 3a for another example). Fig. 1g shows histograms of swimming speeds of cells, before and after adding detergent, indicating that ghosts swim at the same speed as live cells in this, saturating, ATP concentration (*P* = 0.421834 > 0.05 by *t*-test, ratio 0.93 ± 0.24, n = 24, Supplementary Figure 3b). Wild-type ghosts showed a single speed peak around 1.5 μm s^-1^ in detergent (Fig. 1g, bottom), in contrast to peaks at ~1.7 and 3 μm s^-1^ for the same cells without detergent (Supplementary Figure 1b). If CW rotation is associated with the faster peak^17^, and is suppressed by detergent, this is consistent with our lack of observation of CW rotation of beads in the presence of detergent (see below). The lack of the 3 μm s^-1^ peak in cells lacking CheY (Fig. 1g, *top* and Supplementary Figure 4) would then indicate that CheY is required for CW rotation, as in the bacterial flagellar motor^18^. However, we found it difficult to track swimming ghosts due to their low contrast, and were not able to determine the direction of archaellar filament rotation.

To overcome these difficulties, we established a ghost-bead assay for measuring ATP-coupled motor rotation (Fig. 2a). We attached cells with sheared, biotinylated archaellar filaments nonspecifically to the cover glass surface, and then introduced 500 nm streptavidin beads which attached to the filaments (Material and Methods and Supplementary Result 2). Addition of 0.1 mg ml^-1^ streptavidin (which would crosslink adjacent filaments in a rotating bundle) did not stop bead rotation, indicating that shearing removed most filaments and rotating beads are attached to a single archaellum^19^ (Supplementary Video 3). For the preparation of ghosts, live cells labelled with rotating beads were treated in a flow chamber with detergent (0.03 % sodium cholate, as for swimming cells) for less than 30 sec to permeabilize their cell membrane, and subsequently the detergent was replaced with buffer containing ATP. Motor rotation was stopped by permeabilization and reactivated by the addition of ATP (Supplementary Video 4, Fig. 2b). Although beads on ghost cells rotated only CCW in the presence of detergent (n = 11, see Supplementary Result 3 and Supplementary Figure 7), we observed both directions of rotation after detergent removal (Fig. 2c). We did not see any differences between CW and CCW rotation rates (Supplementary Figure 8) and therefore analyzed speeds collectively.

**Figure 2.**
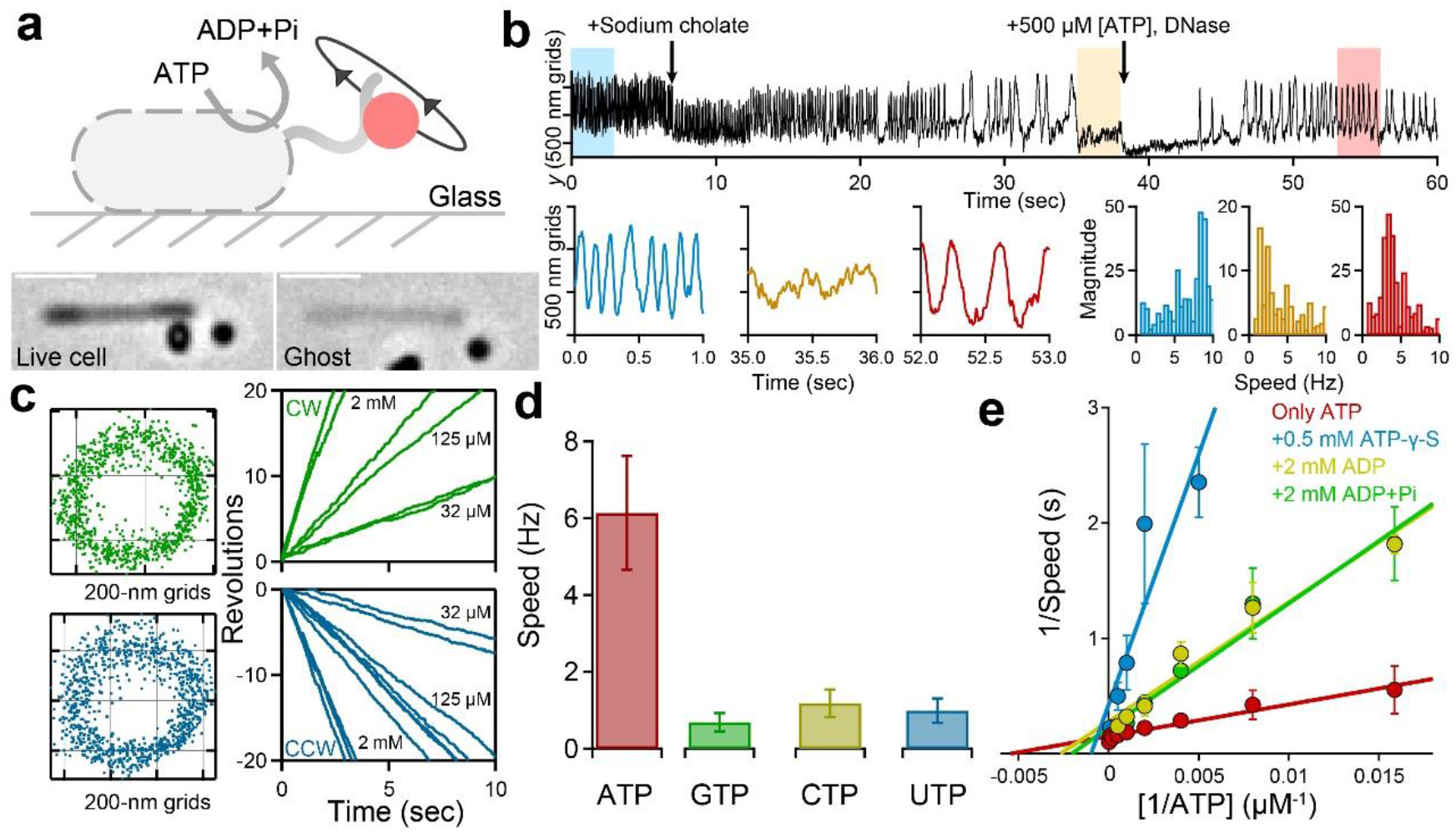
Visualization of motor rotation in ghosts, via beads attached to archaellar filaments. (a) Schematic of experimental setup *(top)* and phase-contrast images of a live cell *(lower left)* becoming a ghost *(lower right).* Scale bar, 5 μm. (b) *Top:* Time course of bead location (*y*-coordinate) during ghost preparation. Bottom: Shaded sections (*top*) are expanded (*left*); with corresponding speed distributions by Fourier transform analysis (*right*), using the same colours as *top*. Blue shows the live cell, orange the motor stopped after treatment with detergent, red the motor re-activated after addition of ATP. Rotation cannot be measured during media exchange time, ~10 s (Data from Supplementary Video 4). (c) *Left: x-y* plots of locations of two different beads attached to archaeallar filaments. Green and blue represent CW and CCW rotation, respectively. *Right:* Angle *vs* time for [the same two/similar (delete whichever is not true)] beads. The slopes decrease in proportion to [ATP] in both directions, indicating ATP-coupled rotation. (d) Rotation rate for different nucleotide triphosphates at 10 mM. The mean ± SD were 6.14 ± 1.48 Hz for ATP (n = 43), 0.69 ± 0.24 Hz for GTP (n = 30), 1.18 ± 0.36 Hz for CTP (n = 32), and 0.99 ± 0.32 Hz for UTP (n = 29). (e) Lineweaver-Burk plot of rotation rate and inhibitors. Blue, green, ocher, and red represent data with 2 mM ADP (n = 140), 2 mM ADP+Pi (n = 114), 0.5 mM ATP-γ-S (n = 118), and without inhibitors (n = 345), respectively. Data are representative of three independent experiments.

We next investigated the effect of different nucleotide triphosphates (NTPs). Previous *in vitro* experiments showed that purified FlaI hydrolyzes different NTPs at similar rates^20^. However, the archaellar rotational rates in ghosts in 10 mM GTP, CTP, and UTP were 5-10 times slower than in ATP (Fig. 2d). This suggests that the motor complex might increase the selectivity of FlaI for nucleotides and/or prevent extra energy consumption *in vivo* like the endopeptidase Clp (see Fig. 1B in Ref. 21). We also tested the inhibitory effects on rotation of ADP, ADP+Pi, and the non-hydrolysable ATP analog ATP-γ-S (adenosine 5’-[γ-thio]triphosphate). We saw no rotation with ATP-γ-S alone. We measured the rotation rates of 500 nm beads attached to archaella in ghosts over a range of [ATP] between 63 μM and 10 mM, with and without each of ADP (2 mM), ADP+Pi (each 2mM) and ATP-γ-S (0.5 mM). Figure 2e shows the results as a Lineweaver-Burk plot. All 3 caused large reduction of rotation rates at lower [ATP], but much smaller reductions of *f*_max_, indicating competitive inhibition. The inhibitor constants, *K*_i_, were estimated to be 1.94 mM for ADP (Ocher), 1.22 mM for ADP.Pi (Green), and 0.11 mM for ATP-γ-S (Blue). We also observed modest effects of pH, and ion concentration on rotation (Supplementary Result 4).

Although we expected bi-directional rotational to be mediated by the response regulator CheY^10^, live cells without CheY were observed to rotate in either direction, without switching during our typical recording time of 30 s (n = 5 for CW rotation, n = 76 for CCW rotation). To observe the role of archaeal CheY in motor switching, we extended our recording time to 300 s. Switching from CCW to CW rotation was frequent in wild type live cells, but rare in ΔCheY live cells even during 5 min recordings (Fig. 3 and Supplementary Figure 10). Wild type ghosts still switched, but the bias and fraction of switching cells were changed, suggesting the chemotaxis system was still active, but altered (Supplementary Table 3).

**Figure 3.**
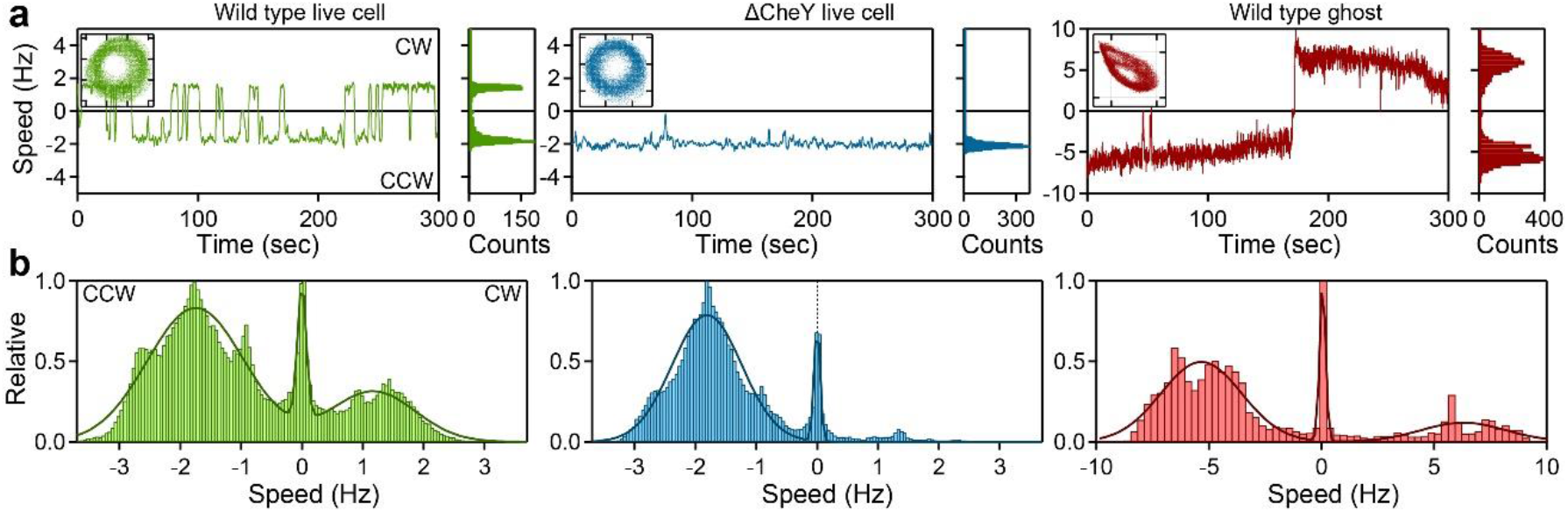
Archaeal CheY-mediated motor switching. (a) *Left:* Representative time course of rotation rate for 5 min, using 970 nm bead in wild type- and ΔCheY live cells, and 500 nm bead in wild type ghosts. Positive and negative speeds represent CW and CCW rotation, respectively. *Inset: y-x* plots of bead rotation. Grids represent 500 nm. *Right:* Histogram of rotation rate of each cell. (b) Population histograms of rotation rate. The solid line represents a Gaussian distribution, where the peak and SD were 1.17 ± 0.31 Hz for CW rotation and 1.75 ± 0.79 Hz for CCW rotation in wild type cells (46 cells); 1.81 ± 0.57 Hz for CCW rotation in the ΔCheY live cells (54 cells); and 6.31 ± 1.90 Hz for CW rotation and 5.34 ± 1.79 Hz for CCW rotation in wild type ghosts (28 cells). Data are representative of two independent experiments.

Figure 4a shows the dependence of rotation speed *f*, revs per second) of 200, 500, and 970 nm beads upon [ATP] in the range 8 μM to 10 mM. Michaelis-Menten fits to the data (solid lines) are poor below 30 μM ATP. Figure 4b shows the relationship between *log([ATP])* and *log(f / (f_max_-f)),* where *f_max_* is estimated by Michaelis-Menten plot (Fig. 4a), and the slope represents Hill coefficients of 0.63, 0.82 and 0.89 for 200 nm, 500 nm, 970 nm beads respectively. This result indicates negative cooperativity in ATP-driven archaellar rotation, (see below for discussion). Figure 4c shows the relationships between torque, speed and [ATP] for archaellar rotation. The maximum motor torque was estimated to be ~200 pN nm for live cells and ~170 pN nm for ghosts, comparable to *Hbt. salinarum* live-cell experiments (160 pN nm)^22^.

**Figure 4.**
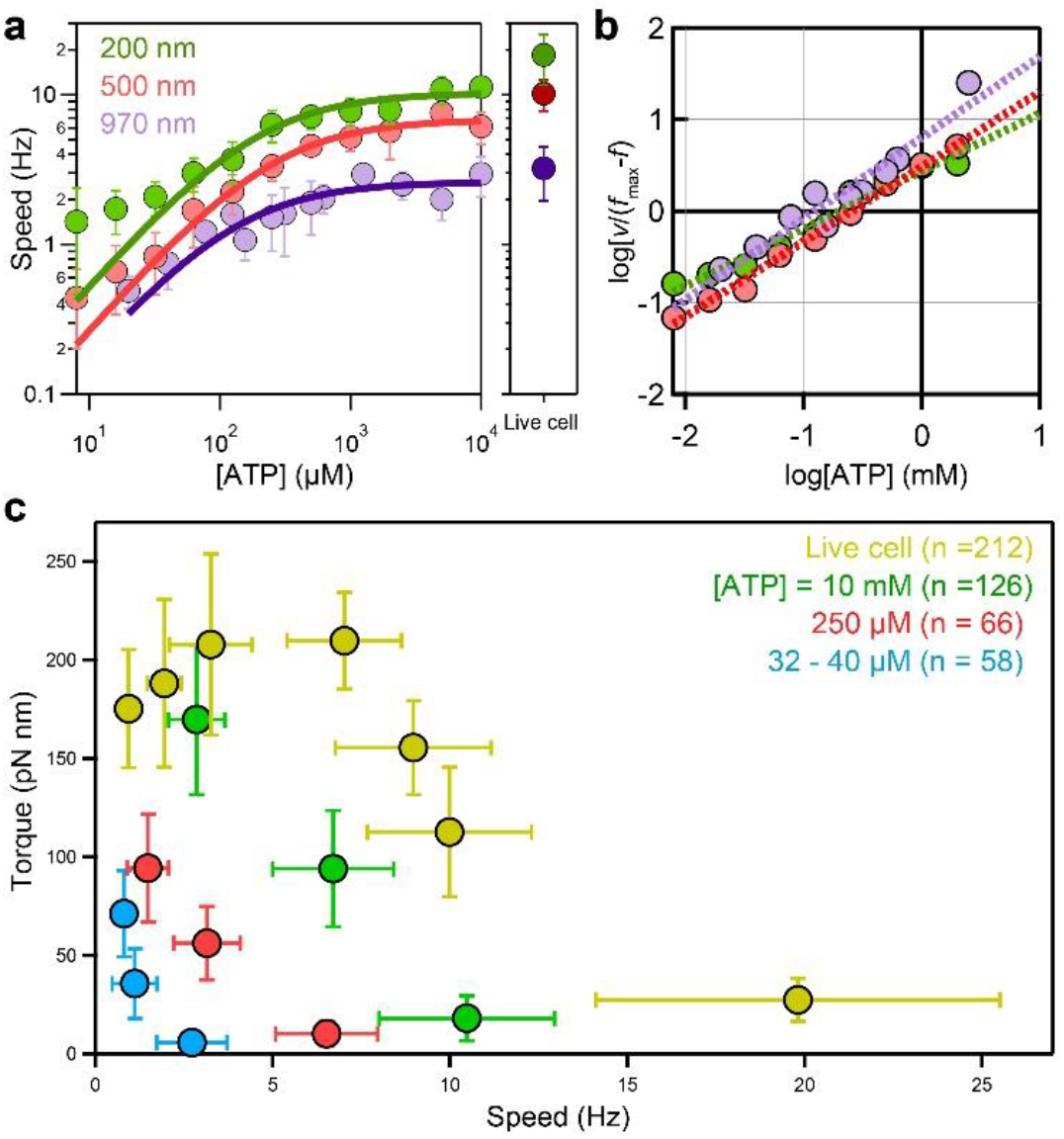
ATP-and load-dependent archaeal motor rotation. (a) Rotation rates of 200 (green), 500 (red) and 970 nm beads (blue) attached to archaellar filaments, *vs* [ATP]. The solid lines show a fit to the Michaelis-Menten equation 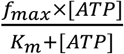; where *f*_max_ and *K*_m_ are 10.3 Hz, 188 μM for 200 nm beads (n = 287), 6.8 Hz and 249 μM for 500 nm beads (n = 438), and 2.6 Hz and 132 μM for 1000 nm beads (n = 303). *Right*: corresponding rotation rates of live cells; 18.47 ± 6.86 Hz for 200 nm beads (n = 19), 10.17 ± 2.39 Hz for 500 nm beads (n = 32) and 3.22 ± 1.27 Hz for 1000 nm beads (n = 72). (b) A Hill plot of the same data. The Hill coefficient, determined from the slope of the plots, was 0.63 for 200 nm beads, 0.82 for 500 nm beads, and 0.89 for 970 nm beads. (c) Torque *vs* speed (mean ± SD), of live cells and ghosts at various [ATP]. Torque was estimated as *T* = 2*πfξ*, where *f* is rotation speed and *ξ* = 8*πηa*^3^+6*πηar*^2^ the viscous drag coefficient of the bead. *ξ* was varied by using bead size (*d* = 2*a* = 200, 500, 970 nm) and viscosity (*η* = 2.5, 3.9, 7.2 mPa·s in buffer, 5% and 10% Ficoll respectively). *r* is the major axis of the ellipse describing the orbit of the bead center. [ATP] and number of cells are indicated.

Our estimated maximum torque of ~170 pN nm corresponds to the motor doing work for a single rotation (2πT, ~1000 pN nm) equivalent to the free energy of hydrolysis of ~12 ATP molecules per revolution, assuming the free energy of 80-90 pN nm per ATP molecule. Conservation of energy therefore sets a lower limit of 12 /rev/motor on the ATP hydrolysis rate, ~15 times higher than that measured *in vitro* for FlaI^23^. This indicates that motor assembly enhances ATPase activity in the archaellum, as observed in other systems; for example the PilC-PilT interaction in T4P^24^ and β- and γ-subunit interaction in F1-ATPase^25^. Hydrolysis of 12 ATP molecules per revolution is consistent with previous reports^22^ and with models of a 2-fold FlaJ rotor rotating within a 6-fold FlaI ATPase^22,26^.

Negative cooperativity in archaellar rotation at low [ATP] (Fig. 4a-b) might be explained by a mechanism similar to that proposed for F1-ATPase, where bi-site and tri-site hydrolysis correspond to nucleotide occupancy of the three catalytic sites alternating between 1 and 2 or between 2 and 3 or three, respectively^27^, and the hydrolysis rate is slower when only 1 site is occupied. In this scenario, negative cooperativity would be the result of the same interactions within the FlaI hexamer that power rotation^22,26^. Negative cooperativity could also arise from communication between FlaI and FlaH rings, similar to inter-ring effects in chaperonins^28^. Although FlaH has only the Walker A motif, ATP binding is known to modulate the interaction between hexametric rings of FlaI and FlaH 16

Our finding that the time-averaged motor torque decreases with increasing speed at low loads (Fig. 4c) differs from a previous report^22^, which assumed constant torque irrespective of viscous load and speed, and explained the observed speed variations by assuming an extra contribution to the viscous drag from an unseen remnant of the filament. The required length of these remnants (ξ,0.8 pN nm s) would be about 4 μm^22^, which seems unlikely given our observation that most filaments are removed by shearing. The curves in figure 4c are qualitatively similar to those reported for the BFM with varying ion-motive force^29^ By contrast, the equivalent data for F1-ATPase correspond to Michaelis-Menten kinetics and torque that decreases linearly with increasing speed (see Fig. 2 in Ref.^30^). Simple models for the torque-speed relationships, similar to those applied to the BFM^29,31,29,31^ and high-resolution detection of steps in rotation^32^ using gold nanoparticles^30,33^ may reveal the details of the rotation mechanism of the archaellum in future.

So far, there is no direct evidence as to which components of the archaellum are fixed relative to the cell (“stator”) and which rotate with the filament (“rotor”). FlaF and G are most likely part of the stator, anchored to the S-layer. Previous reports indicate interaction between FlaF and the S-layer and a deficiency in swimming motility of S-layer deleted cells^14,15^. Our observations of increased motility with [CaCl_2_] and the speed fluctuations at low [CaCl_2_] (Supplementary Figures 1 and 9), given that calcium stabilizes S-layer^34^, support this hypothesis. Homology to F1-ATPase and T4P are generally taken to favour a model where rotation of a FlaJ dimer within the central core of the FlaI hexamer is driven by cyclic changes in the conformation of the FlaI hexameric ATPase, coupled to ATP hydrolysis^22,26^. In this model, FlaJ is the rotor and all other motor components are the stator (Figure 1b, left). For switching, changes caused by CheY binding, presumably somewhere on FlaC/D/E, would have to propagate all the way to the core of FlaI, which would need separate mechanochemical cycles for CW and CCW rotation. Figure 1b, right, illustrates the other extreme possibility, most similar to the BFM. In this model, conformational changes in FlaI would push on FlaF/G, either directly or via FlaC/D/E. In the latter case, the switch mechanism could reside within FlaC/D/E and FlaI need not have separate modes for CW and CCW rotation. Intermediate models are also possible (Figure 1b, middle). Our ghost model may allow labelling of archaellar components to observe directly which are part of the rotor^35,36^, analysis of rotational steps as in isolated F1 and other molecular motors^30,37,38^, and direct investigations of the role of CheY.

Our finding that archaeal CheY increases CW bias (Fig. 3) is inconsistent with previous reports^10^ that the role of archaeal CheY in *Hbt. salinarum* enhances switching from CW to CCW rotation, as revealed by a dark-field microscopy of swimming cells^10,39^. This study measured static filaments to be right-handed in *Hbt. salinarum* M175, and inferred from this that CW rotation propels a cell forward, CCW backwards. We speculate that the contradictory result might be due to misinterpretation of filament helicity caused by errors in accounting for reflections in the microscope optics - cryo-EM data show left-handed helicity in *Hbt. salinarum* M175, supporting our conclusion^40^. With due care to account for reflections in our microscope (Supplementary figure 6), our bead assay is a direct observation of the rotational direction of the motor.

Our ghost assay represents the first experimental system that allows manipulation of the thermodynamic driving force for an archaeal molecular motor, following previous examples including eukaryotic linear motors^41^, the PomAB-type BFM^38^, and *Mycoplasma* gliding motor^9^. We anticipate that this assay will be helpful for other biological systems. Archaea display chemotactic and cell division machinery acquired by horizontal gene transfer from bacteri^a11,12^. Although the archaellum and bacterial flagellum are completely different motility systems, they share common chemotactic proteins. Theoretically, only our ghost technique allows monitoring of the effect of purified CheY isolated from different hosts on motor switching. Similarly, our ghost cells offer the potential to manipulate and study the archaeal cell division machinery as with *in vitro* ghost models of *Schizosaccharomyces pombe*^42^ Ghost archaea offer the advantages of both *in vivo* and *vitro* experimental methods and will allow the exploration of the universality, diversity, and evolution of biomolecules in microorganisms.

## Material and Methods

### Strain and Cultivation

Strains, plasmids, and primers are summarized in Supplementary Table 1, 2. *Haloferax. volcanii (HfX. volcanii)* cells were grown at 42°C on a modified 1.5 % Ca agar plate (2.0 M NaCl, 0.17 M Na_2_SO_4_, 0.18 M MgCl_2_, 0.06 M KCl, 0.5% (wt/vol) casamino acid, 0.002% (wt/vol) biotin, 0.005% (wt/vol) thiamine hydrochloride, 0.01% (wt/vol) L-tryptophane, 0.01% (wt/vol) uracil, 10 mM HEPES-NaOH (pH 6.8) and 1.5% (wt/vol) Agar). Note that 20 mM CaCl_2_ should be added (Supplementary Result 1). Colonies were scratched by the tip of a micropipette and subsequently suspended in 5ml of Ca liquid medium. After 3h incubation at 37°C, the culture was centrifuged at 5,000 r.p.m and concentrated to 100 times volume. The 20 μl culture was poured into 25-ml fresh Ca medium and again grown for 21 h with shaking of 200 r.p.m at 40°C. The final of an optical density would be around 0.07.

Gene manipulation based on selection with uracil in *ΔpyrE2* strains was carried out with PEG 600, as described previously^43^. For the creation of KO strains, plasmids based on pTA131 were used carrying a *pyrE2* cassette in addition to ~1000-bp flanking regions of the targeted gene. *flgA1(A124C)* was expressed by tryptophane promotor (Supplementary Result 2).

### Preparation of biotinylated cells

The culture of *HfX. volcanii* Cys mutant was centrifuged and suspended into buffer A (1.5 M KCl, 1 M MgCl_2_, 10 mM HEPES-NaOH pH 7.0). Cells were chemically modified with 1 mg ml^-1^ biotin-PEG2-maleimide (Thermo Fischer) for 1 h at room temperature, and excess biotin was removed with 5,000 g centrifugation at R.T for 4 min.

### Motility assay on soft-agar plate

A single colony was inoculated on a 0.25% (wt/vol) Ca-agar plate and incubated at 37°C for 3-5 days. Images were taken with a digital camera (EOS kiss X7; Canon).

### Microscopy

All experiments were carried under an upright microscope (Eclipse Ci; Nikon) equipped with a 40× objective (EC Plan-Neofluar 40 with Ph and 0.75 N.A.; Nikon) or 100× objective, a CMOS camera (LRH1540; Digimo). Images were recorded at 100 fps for 10-30 sec. For a motility experiment at 45°C, a phase-contrast microscope (Axio Observer; Zeiss) equipped with a 40× objective (EC Plan-Neofluar 40 with Ph and 0.75 N.A.; Zeiss), a CMOS camera (H1540; Digimo), and an optical table (Vision Isolation; Newport) were used.

For fluorescent experiment, a fluorescent microscope (Nikon Eclipse Ti; Nikon) equipped with a 100× objective (CFI Plan Apo 100 with Ph and 1.45 N.A.; Nikon), a laser (Nikon D=eclipse C1), an EMCCD camera (ixon^+^ DU897; Andor), and an optical table (Newport) were used. The dichroic mirror and emitter were Z532RDC (C104891, Chroma) and 89006-ET-ECFP/EYFP/mcherry (Chroma) for an FM4-64 experiment, Z442RDC (C104887, Chroma) and 89006-ET-ECFP/EYFP/mcherry for an S-layer experiment, and Z442RDC and ET525/50m (Chroma) for a Dylight488 experiment.

### Construction of swimming ghosts

The flow chamber was composed of a 22×22 coverslip and slide glass. Two pieces of double-sided tape, cut to a length of ~30 mm, were used as spacers between coverslips^17^ Two tapes were fixed with a ~5 mm interval, and the final volume was ~15 μl. The glass surface was modified with a Ca medium containing 5 mg ml^-1^ bovine serum albumin (BSA) to avoid cells attaching to a glass surface.

To construct swimming ghosts, 10 ul of cell culture in Ca medium and buffer B (2.4 M KCl, 0.5 M NaCl, 0.2 M MgCl_2_, 0.1 M CaCl_2_, 10 mM HEPES-NaOH pH 7.2) containing 1 mg ml^-1^ DNase, 5 mM ATP (A2383, Sigma Aldrich), and 0.03 % sodium cholate (Sigma Aldrich) was mixed in an Eppendorf tube. Subsequently, the 20 ul mixture was infused into the flow chamber.

Phase-contrast images were captured at 20 frames s^-1^ for 15 sec. Swimming trajectories were determined by the centroid positions of cells and subjected to analysis using Igor pro. Given the trajectory of cells, ***r***(***t***) = [*x*(*t*), *y*(*t*)], the swimming velocity ***v(t)*** was defined as 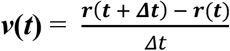.

### Bead assay

For the observation of a rotational bead attached to an archaellar filament, archaellar filaments were sheared by 30 times pipetting with 200 μl pipette (F123601, Gilson), infused into a flow chamber and kept for 10 min. Streptavidin-conjugated fluorescent beads (200 nm (F6774, Molecular probes), 500 nm (18720, Polysciences) or 970 nm (PMC 1N, Bangs lab)) in buffer B (2.4 M KCl, 0.5 M NaCl, 0.2 M MgCl_2_, 0.1 M CaCl_2_ 10 mM HEPES-NaOH pH 7.2, 0.5 mg ml^-1^ BSA (Sigma Aldrich)) were added into the flow chamber, incubated for 15 min, and then rinsed with buffer to remove unbound beads. The solution was replaced to buffer B containing 0.03 % sodium cholate hydrate (C1254, Sigma Aldrich) and 1 mg ml^-1^ DNase. When the optical density of cells was decreased, buffer B was replaced with buffer A containing ATP. Rotary ghosts were prepared within 1 min (Fig. 2b). For pH measurements, the following buffer was used: Bis-Tris HCl for pH5.7 and 6.1 experiments; HEPES-NaOH for a pH8.0 experiment; and Tris-HCl for pH8.6 and pH9.3 experiments (Supplementary Figure 9). For nucleotides experiments, ADP (A2754, Sigma Aldrich), ATP-γ-S (A1388, Sigma Aldrich), GTP (ab146528, Abcam), CTP (R0451, Thermo Fischer Scientific), and UTP (R0471, Thermo Fischer Scientific) were used. Most data were collected at 100 frames s^-1^ for 10 sec.

Bead position was determined by centroid fitting, giving cell trajectories, ***r(t)*** = [*x*(*t*), *y*(*t*)]. The rotation rate was determined from either Fourier transform analysis (Fig. 2b) or a fit with a linear function to time course of bead rotation. (Fig. 2c). The rotational torque against viscous drag was estimated as *T* = 2*πfξ*, where *f* is rotational speed and = 8*πηa*^3^+6*πηar*^2^ the viscous drag coefficient, with *r* the radius of rotation (major axis of ellipse), *a* the bead radius, and *η* the viscosity. We neglected the viscous drag of filaments^22^, which is expected to be negligible compared to these beads^44^.

To measure the viscosity of the medium, we tracked diffusing fluorescent beads for 30 sec at 100 fps and performed an analysis of their mean-squared displacement versus time. From this analysis, the viscosities are estimated to be 0.0025 Pa·s in buffer, 0.0039 Pa·s in buffer + ficoll 5 %, and 0.0072 Pa·s in buffer + ficoll 10 % at 25 °C, which are slightly higher than a previous estimate^22^. We inferred that this discrepancy might be due to the proximity of the glass surface^45^.

### Fluorescent experiments

For visualization of archaellar filaments, biotinylated cells were subsequently incubated with 0.1 mg ml^-1^ Dylight488-streptavidin (21832, Invitrogen) for 3 min, washed by centrifugation, and resuspended.

FM4-64 (F34653, Life Sciences) was used to stain the archaeal cell membrane. The powder was dissolved by buffer B (1.5 M KCl, 1 M MgCl_2_, 10 mM HEPES-NaOH pH 7.0), and the cells were incubated for 30 min. The extra dye was removed by centrifugation. For microscopic measurements, the glass surface was cleaned using a plasma cleaner (PDC-002; Harrick plasm).

Quantum dots 605 (Q10101MP, Invitrogen) was used to stain the archaeal cell surface, S-layer^17^ Cells were biotinylated with biotin-NHS-ester (21330, ThermoFisher) and incubated for 15 min at R.T. Extra biotin was washed by centrifugation. Biotinylated cells were subsequently incubated with the buffer containing QD605 at a molar ratio of 400:1 for 3 min, washed by centrifugation, and resuspended.

## Author Contributions

Y.K. and R.M.B designed the research. Y.K. performed all experiments and obtained all data; N.M. helped genetics, biochemistry, and preparation of figures; Z.L, F.B., T.EF.Q., C.v.d.D and S.-V. A. helped genetics; R.I helped the ghost experiments; N.M. and R.M.B helped microscope measurements; Y.K., and R.M.B. wrote the paper.

## Acknowledgements

We thank Prof. Achillefs Kapanidis and Dr. Abhishek Mazumder for sharing chemicals, Dr. Nariya Uchida for sharing his useful information in the torque calculation, and Dr. Mitsuhiro Sugawa for the technical advice in the microscope measurement. This study was supported in part by a grant from the Funding Program for the Biotechnology and Biological Sciences Research Council (to R.M.B), Collaborative Research Center Grant from the Deutsche Forschungsgemeinschaft (to S-V.A.). Y.K was recipient of the Japan Society for the Promotion of Science Postdoctoral Fellowship for Research Abroad and the Uehara Memorial Foundation postdoctoral fellow, and N.M. was recipient of the Yoshida Scholarship Foundation.

