## Supplementary Information for "Motile ghosts of the halophilic archaeon, *Haloferax volcanii*"

#### This file includes:

Supplementary Results 1-5

Supplementary Figures 1-11

Supplementary Tables 1-3

Captions for Supplementary Videos

Supplementary References

### Supplementary Results

#### 1. Swimming motility of *Haloferax volcanii*

*Haloferax volcanii* (*Hfx. volcanii*), a halophilic archaeon, requires more than 3M salt for growth and is one of the most genetically abundant species [1,2]. Under an optimal-growth condition, a single cultivated cell is rod shaped with a diameter of 0.9 μm and length of 2-10 μm, and swims with 180°-reversals. We increased the fraction of swimming *Hfx. volcanii* cells from 20-30 % [2] to 80 % by adding 20 mM $CaCl_2$ (Supplementary Video 1 and Supplementary Figure 1a). The distribution of swimming speeds has two peaks, at $v_{slow} = 1.72 \pm 0.25$ μm $s^{-1}$ and $v_{fast} = 2.95 \pm 0.60$ μm $s^{-1}$ at room temperature (n = 72), $v_{slow} = 2.44 \pm 0.27$ μm $s^{-1}$ and $v_{fast} = 3.87 \pm 0.88$ μm $s^{-1}$ at 45°C (n = 115, Supplementary Figure 1b). A previous study reported that CW rotation produces a thrust 1.5-fold more efficient (*v* swimming speed/ *f* rotation rate) compared to that of CCW rotation [3,4], suggesting that the slow and fast peaks correspond to CCW and CW rotation, respectively. The 20% increase in swimming speed at 45 °C compared to 25°C, is similar to previous results for a different species of halophilic archaeon, *Halobacterium* (*Hbt.*) *salinarum* [3]. All experiments were performed at room temperature to facilitate preparation of ghosts.

#### 2. No effect of cysteine mutation in filaments on swimming motility

We previously showed that the fluorescent markers could be attached to archaellar filaments through a biotin-avidin interaction [3,5,6], but this method is not efficient for *Hfx. volacnii*. To effectively attach a streptavidin bead to an archaellar filament, we genetically introduced cysteine substitutions into the environment-exposed surface of the archaellins FlgA1(A124C) (Supplementary Figure 5a). The expression level of FlgA1(A124C) was controlled by tryptophan in a Δ*flgA1* strain, and motility was restored with 1 mM [tryptophan], but not in the absence of tryptophan (Supplementary Figure 5b). We could not see any difference in diameter on a semi-solid agar plate (Supplementary Figure 5c) and in speed in solution, where the average speed and SD were 1.65 ± 0.14 μm $s^{-1}$ and 2.72 ± 0.49 μm $s^{-1}$ in FlgA1(A124C) cells (n = 75, Supplementary Figure 5d).

**3. The effect of detergent on rotation in bead assay**

We optimized the experimental condition to minimize the effect on motor rotation of the detergent used to prepare ghosts, using 500 nm beads. We tested different types of detergent over the range 0.01-0.1%: Triton X-100, Tween 20, CHAPS, and *n*-Dodecyl-β-D-maltopyranoside (DDM) and sodium cholate. With Triton X-100, Tween 20, and CHAPS, the cells were burst and vanished. Cells were mildly permeabilized using DDM or sodium cholate hydrate; however with DDM their shape changed from rod to sphere (data not shown) and very few beads were seen to rotate, with an average speed of of 4.58 ± 1.76 Hz at 2 mM [ATP] (n = 4). Therefore we selected sodium cholate for further optimization. At 0.015 mM, ~5 minutes exposure was required before ghosts were seen, whereas ~10 s was sufficient to produce ghosts at 0.03 and 0.05 %. The rotational rate of ghosts was the fastest with 0.03% (Supplementary Figure 7a). We inferred that the motor complex might be damaged in proportion to exposure time to detergent, as shown in Supplementary Figure 7b,c. Taken together, we chose ~10 s exposure to 0.03 % sodium cholate hydrate as the best condition for ghost preparation.

**4. Characterization of other dependences of bead rotation**

We examined the effect of pH on the rotational rate of both ghosts and live cells (Supplementary Figure 9). We detected the rotation on ghosts over the range pH 5.7 to 9.3, but none at pH 5.3. Live cells also rotated at pH 5.3. The rotation speed in ghosts was independent of pH, while that in live cells declined with increasing pH. We also examined the effects of $K^+$ and $Ca^{2+}$ (Supplementary Figure 9a, b respectively). Both ghosts and live cells slowed slightly as [KCl] increased from 1 M to 3 M, the effect stronger in ghosts (Supplementary Figure 9b). Rotation speed and stability were much reduced below ~10 mM [$CaCl_2$]. At higher [$CaCl_2$], speed reduced slightly but stability increased (Supplementary Figure 9c). Slow rotation and frequent pauses could be observed at low [$CaCl_2$], while most ghosts exhibited smooth rotation with a speed of approximately 6 Hz at 100 mM [$CaCl_2$]. Because calcium is essential for maintaining the S-layer [7], this result suggested that the interaction between the stator (FlaF) and the S-layer became weak and/or abolished at low [$CaCl_2$], and consequently, the open-close conformational changes of the FlaI cylinder might not be efficient for generating rotation [8].

**5. Switching behavior of ghosts**

No switching behavior was detected in over 3000 ghosts with observations for 10 to 30 sec. Long-term observations of rotation for 5 min revealed motor switching (Fig. 3a). Of the population of wild type ghosts 24 % rotated exclusively CW during 5-mins recording, 66 % exclusively CCW, and 10 % changed their rotation during the recording (Supplementary Table 3); the CW bias was estimated to be 0.27 (n = 29).

Why do ghosts exhibit so little motor switching? The archaeal chemotactic system is probably related to the CheC-CheD-CheY system in *Bacillus subtilis* [9]*. In B. subtilis*, the histidine kinase CheA autophosphorylates by sensing chemotactic signals on Methyl-accepting Chemotaxis Proteins [10,11], and phosphorylates the response regulator CheY. CheY-P regulates motor switching, and then is modulated by two phosphatases CheC or FliY, instead of CheZ like *E. coli* system. A phosphatase activity of FliY is constitutively on, whereas that of CheC is regulated by CheD. Because the constant CW rotation was detected in ghosts (Supplementary Table 3), we speculated that low switching behavior might be due to lack of phosphatase CheC. Indeed, *B. subtilis* reports showed that ΔCheC mutant does not change the rotational bias but reduces the switching frequency, which is consistent with our observation [12,13]. Another possibility is loss of CheA activity or lack of CheY in solution and consequently not enough for motor switching. More experimental results should be needed for more discussions.

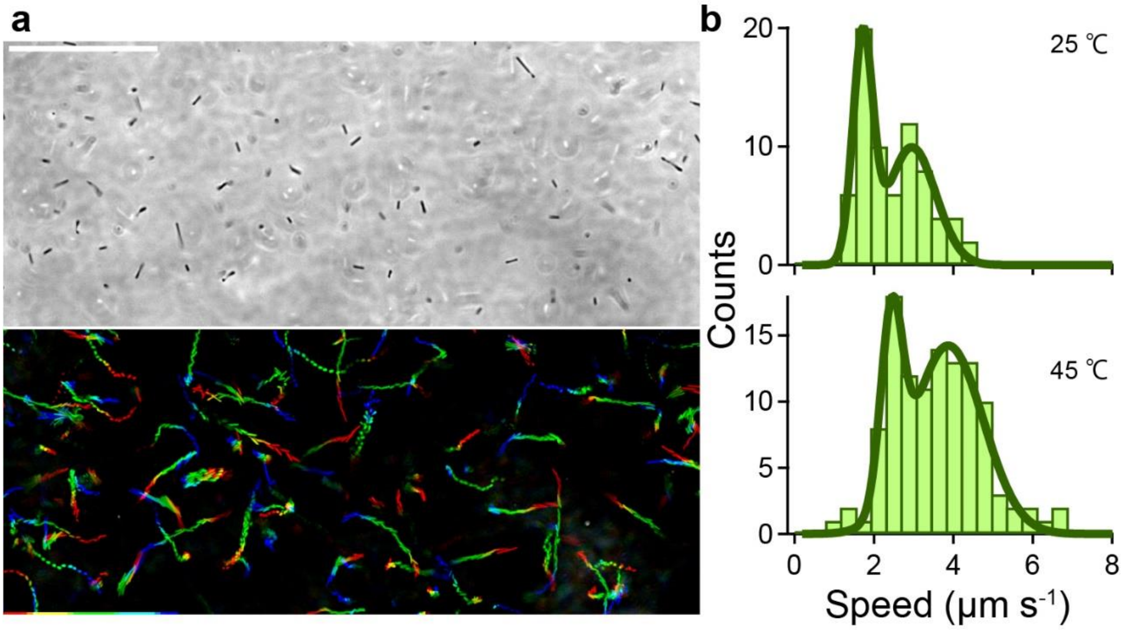

**Supplementary Figure 1 Swimming motility of *Hfx. volcanii* live cells.**

(a) *Top*: Phase-contrast image of *Hfx. volcanii* wild type, FYK1. Scale bar, 50 μm. *Bottom*: Sequential phase-contrast images with 500-ms intervals were integrated for 15 s with the intermittent color code "red → yellow → green → cyan → blue." (b) Histograms of swimming speed at room temperature, R.T. (n = 72) and 45 °C (n = 115), respectively. Solid line represents the Gaussian distribution with 1.72 ± 0.25 μm $s^{-1}$ and 2.95 ± 0.60 μm $s^{-1}$ at R.T. and 2.44 ± 0.27 μm $s^{-1}$ and 3.87 ± 0.88 μm $s^{-1}$ at 45°C, respectively ($P = 5.873\times10^{-10} < 0.05$ by *t*-test). Data are representative of at least two independent experiments.

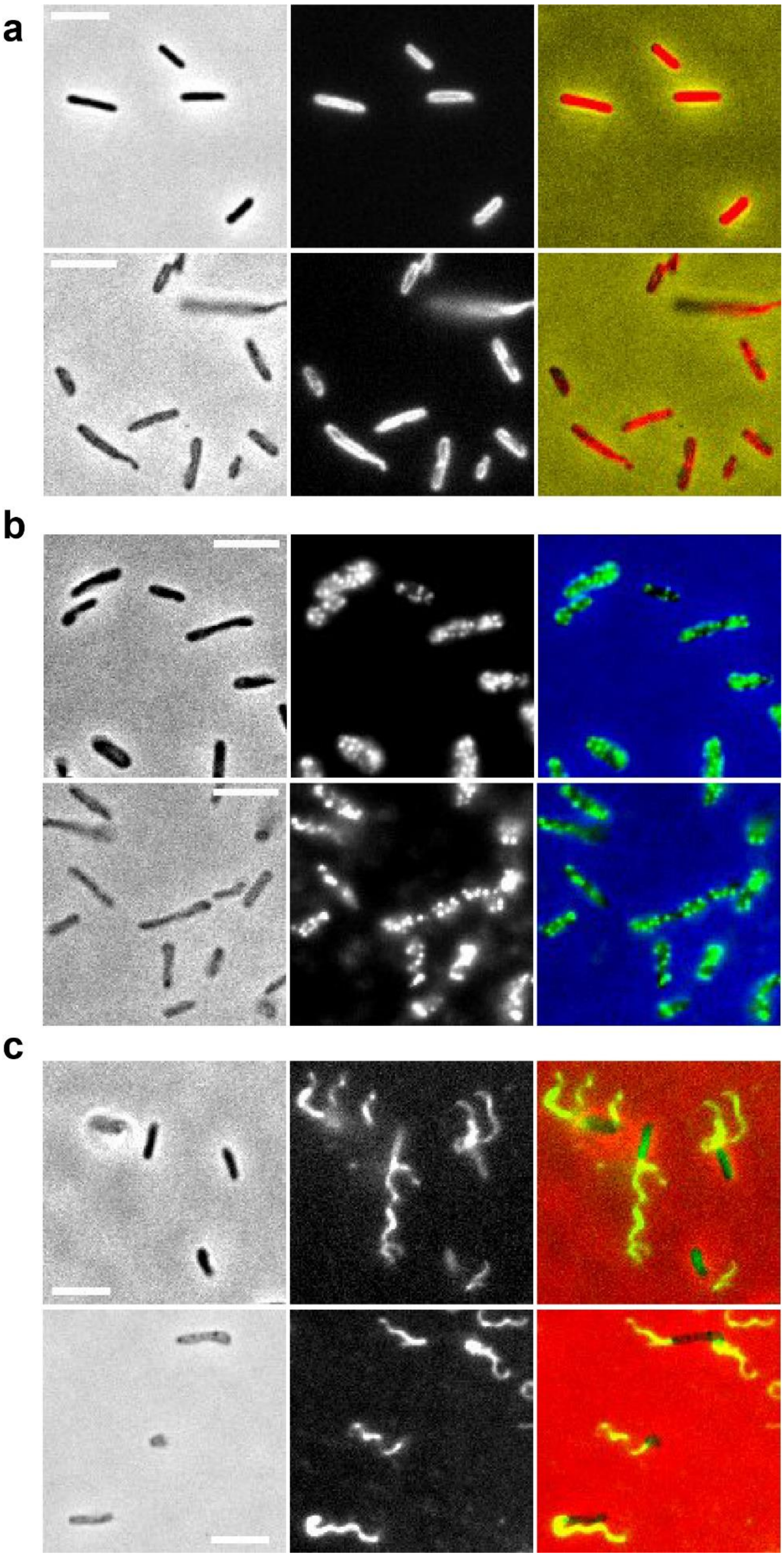

106

#### Supplementary Figure 2 Visualization of filaments and cell membrane

(a) Cell membrane of live cell (*top*) and ghost (*bottom*) fluorescent stained with FM4-64. *Left*: Phase-contrast image. *Middle*: Fluorescent image. *Right*: Merge. The pseudo color of phase-contrast and fluorescent images were red and yellow, respectively. (b) S-layer of live cell (*top*) and ghost (*bottom*) fluorescent stained with Quantum dots 605. *Left*: Phase-contrast image. *Middle*: Fluorescent image. *Right*: Merge. The pseudo color of phase-contrast and fluorescent images were blue and green, respectively. (c) Archaellar filaments of live cell (*top*) and ghost (*bottom*) stained with dylight488. *Left*: Phase-contrast image. *Middle*: Fluorescent image. *Right*: Merge. The pseudo color of phase-contrast and fluorescent images were red and green, respectively. These phenomena were observed in two independent experiments. Scale bar, 5 μm. Data are representative of at least two independent experiments.

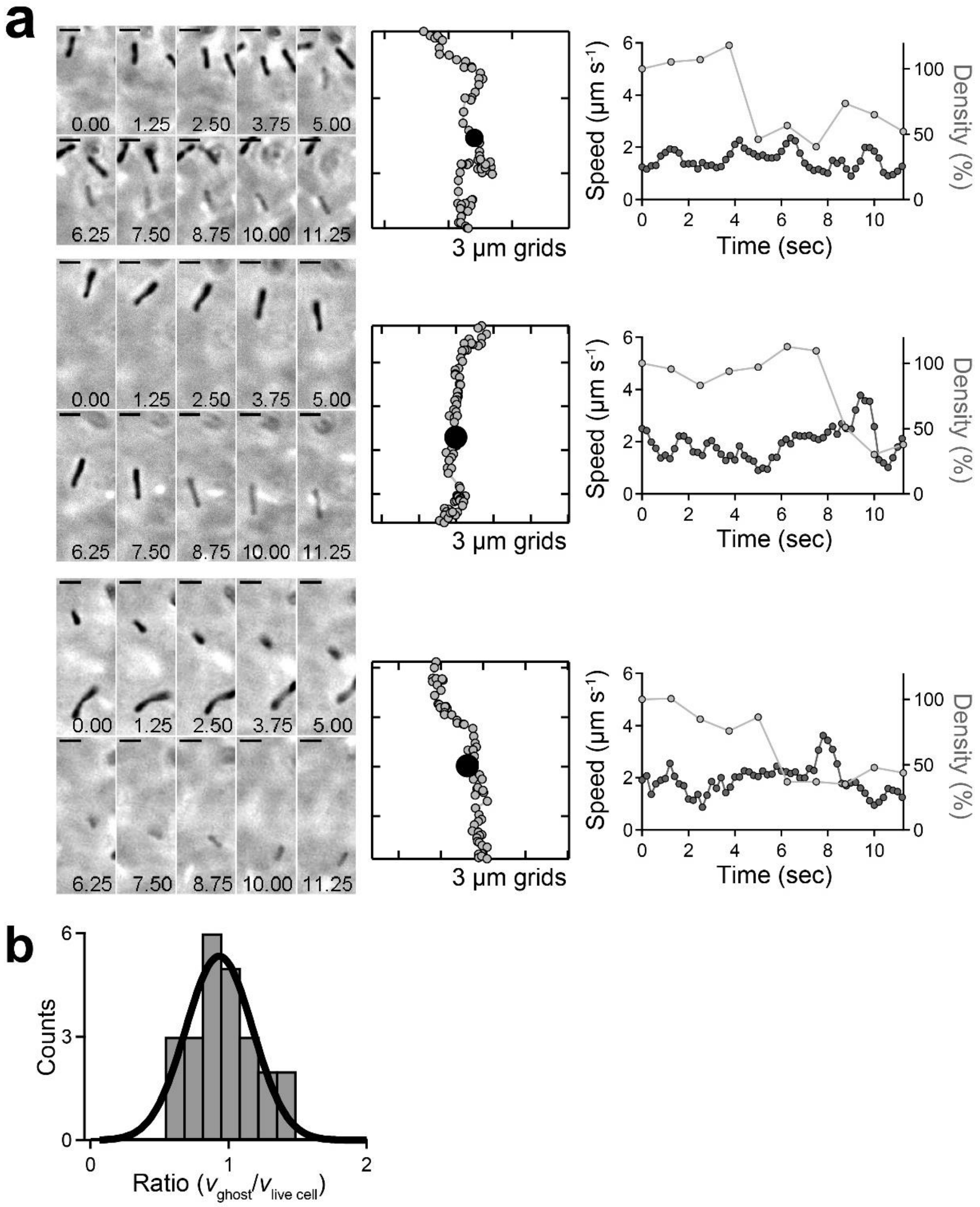

**Supplementary Figure 3 Examples of swimming ghosts.**

(a) *Left*: Sequential phase-contrast images of changes from a live cell to ghost in a free-swimming state at 1.25 s-intervals. Scale bar, 5 µm. *Middle*: Swimming trajectories. Each dot represents one cell localization, at 0.2-sec intervals. Live cells change into ghosts at the black dots. *Right*: Time course of a swimming speed ($v$, black) and cell density (gray). (b) Histogram of the ratio between $v_{live\ cell}$ and $v_{ghost}$. The peak and SD were 0.93 ± 0.24. (n = 24). Data are representative of nine independent experiments.

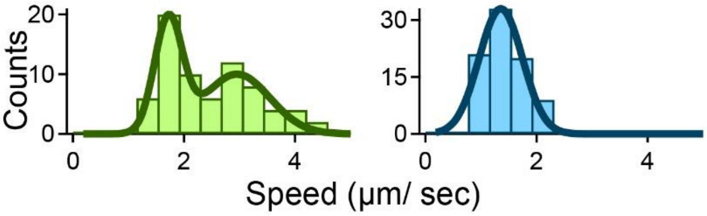

**Supplementary Figure 4 Swimming motility of ΔCheY live cells**

Histograms of swimming speed in wild type (*left*) and ΔCheY (*right*). ΔCheY cells, unlike wild type cells, had a single peak in a swimming speed in liquid media, where the average and SD were 1.36 ± 0.39 µm $s^{-1}$ (n = 84). Considering that CW rotation produces a 1.5-fold more efficient thrust compared to that of CCW rotation [3,6,14], the archaellum motor probably rotates CCW in the absence of the response regulator CheY. The wild-type data comes from Supplementary figure S1b *top*.

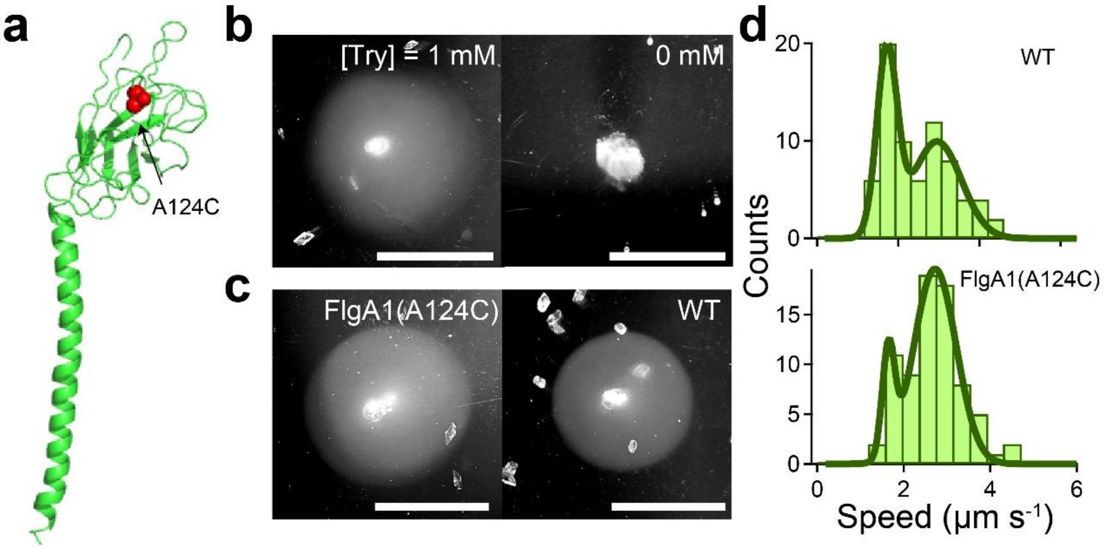

**Supplementary Figure 5 No effect of cysteine substitution in archaellar filament on motility.**

(a) Predicted structure of the archaellar filament, FlgA1, in *Haloferax volcanii* (calculated by Phyre V 2.0; illustration created with PyMOL). The alanine 124 is substituted to cysteine (red sphere). (b) Motilities of *Hfx. volcanii* FlgA1(A124C) cells (FYK3) on 0.25% (wt/vol) soft-agar plates at 37ºC after 3-5 days incubation with induction of 1 mM tryptophane (*left*) and without tryptophane (*right*). Scale bar, 1 cm. (*c*) Motilities of FYK1 (wild type) and FYK3 cells on 0.25% (wt/vol) soft-agar plates at 37ºC after 3-5 days incubation. The diameter of the colony was almost the same, indicating no influence of cysteine substitution on motility. Scale bar, 1 cm. (*d*) Histograms of swimming speed of FYK1 (top, n= 72) and FYK3 (bottom, n = 75) at room temperature. Solid line represents the multiple Gaussian distribution with 1.72 ± 0.25 μm $s^{-1}$ and 2.95 ± 0.60 μm $s^{-1}$ at FYK1 and 1.65 ± 0.14 μm $s^{-1}$ and 2.72 ± 0.49 μm $s^{-1}$ FYK3, respectively ($P$ = 0.1033> 0.05 by *welch's t*-test). These phenomena were observed by at least two independent experiments. Data are representative of at least two independent experiments.

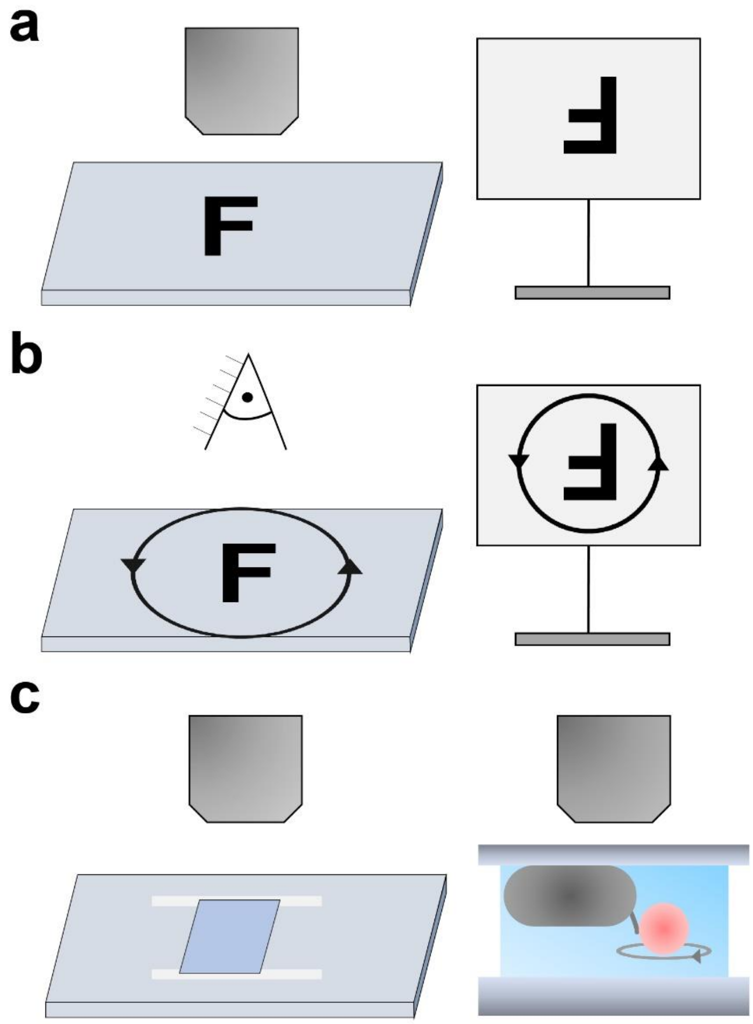

**Supplementary Figure 6 Determination of rotational direction of motor**

(a) A schematic of how we determined the rotational direction. In this study, we used an upright microscope and captured images through a high-speed CMOS camera. The orientation and chirality of a sample in the lab are represented on the left, the corresponding digitized image on the right. Our images are rotated by not reflected (b) Rotation direction in the digitized image is the same as viewed from above (objective side) in the lab. We verified this by observing manual rotation of a microscope slide in the CCW direction as viewed from above - the image also rotated in CCW direction. (c) (c) A cell shows CCW rotation of beads attached to short archaellar filaments as viewed looking along the filament towards the cell, following the convention used for the BFM [15]. In our experiments, cells are attached to the top coverglass, and CCW rotation of beads corresponds to CW rotation in the digitized image. We call this CCW rotation.

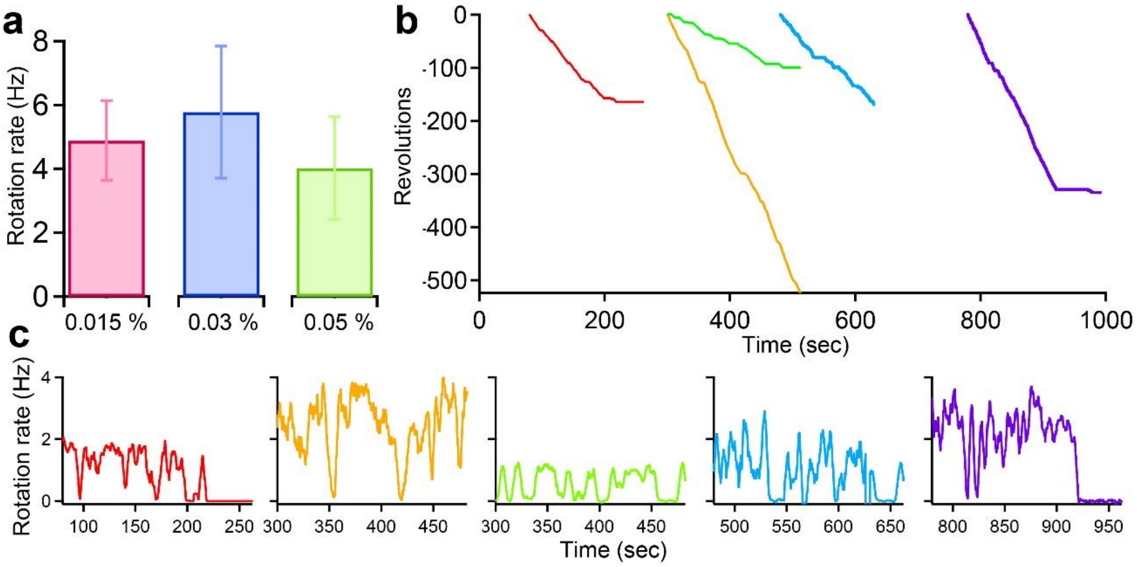

**Supplementary Figure 7 Effect of [detergent] on rotation.**

(*a*) Effect of [detergent] on rotation at 2mM [ATP]. The average and SD were 4.88 ± 1.25 Hz at 0.015% sodium cholate hydrate (n = 33), 5.78 ± 2.07 Hz at 0.03 % (n = 58), and 4.02 ± 1.61 Hz at 0.05% (n = 22). (b) Time course of revolutions in the presence of 0.03% sodium cholate hydrate and 2 mM ATP. Their colors coincide with c. (c) Time course of speed. Motor rotation frequently and often permanently stopped in the presence of detergent. Data are representative of at least two independent experiments.

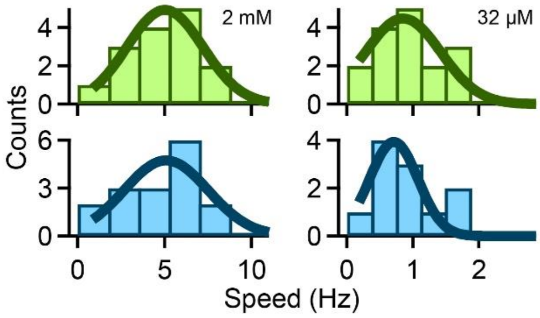

**Supplementary Figure 8 Bidirectional rotation at various [ATP]**

The rotation rates were 0.85 ± 0.57 Hz in CW (*top*, n = 16) and 0.71 ± 0.37 Hz in CCW rotations (*bottom*, n = 12) at 32 μM [ATP] (*P* = 0.4003 >0.05 by *t*-test), and 5.02 ± 2.18 Hz in CW (n = 15) and 5.09 ± 2.41 Hz in CCW rotations (n = 16) at 2 mM [ATP] (*P* = 0.9801 >0.05 by *t*-test).

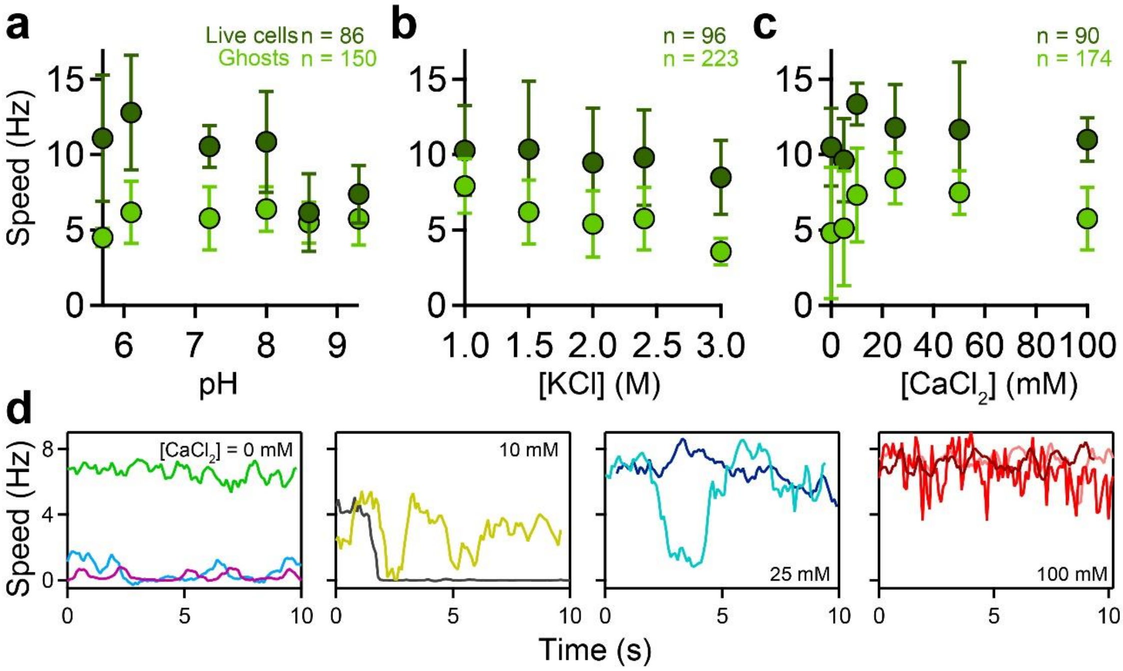

**Supplementary Figure 9 Effect of [ion] and pH on rotation.**

(*a-c*) Effect on rotation rate of live cells (dark green) and ghosts in 2 mM ATP (light green): of pH (a); [KCl] (b); [$CaCl_2$] (c). (d) Time course of speed. The rotation rate was frequently changed and slow at low [$CaCl_2$]. Data are representative of at least two independent experiments.

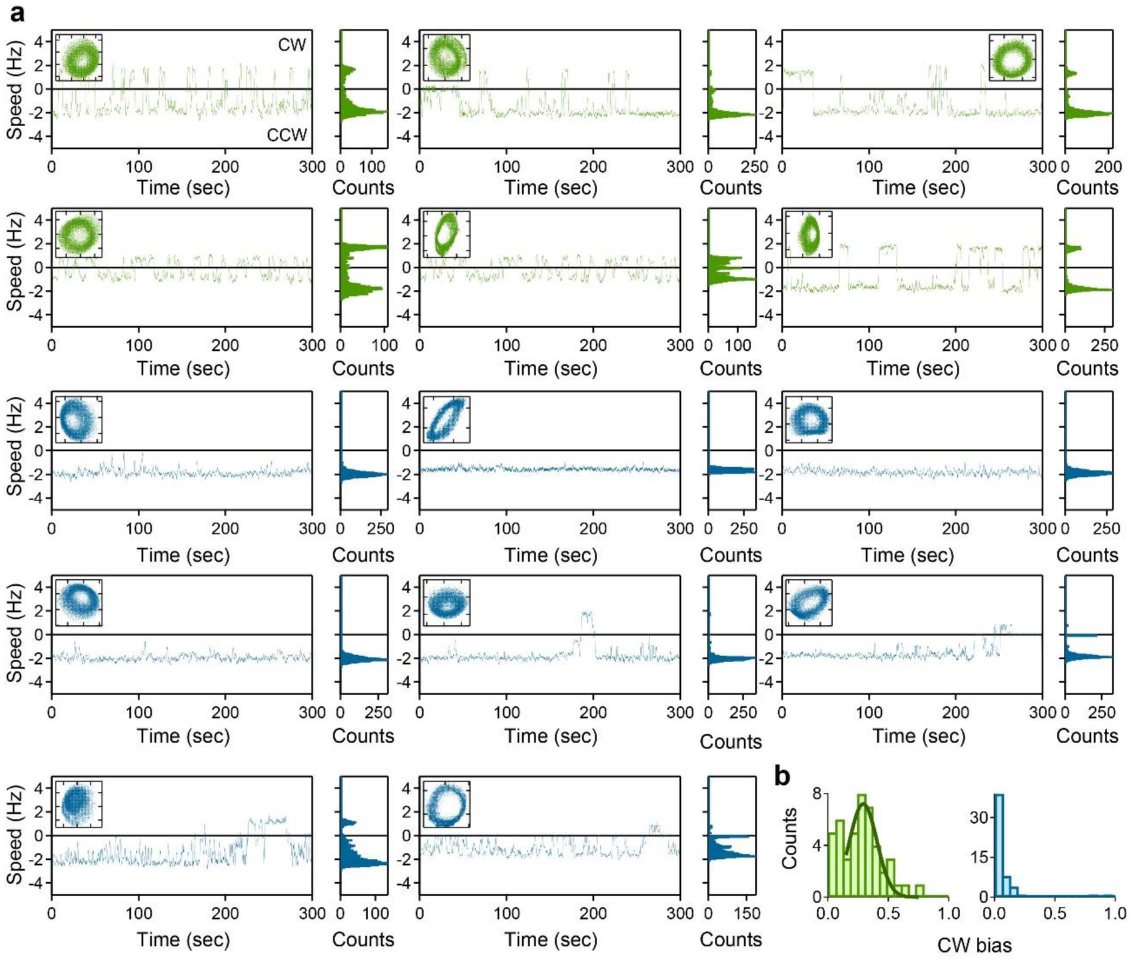

**Supplementary Figure 10 Long-term observation of motor switching**

(a) *Left*: Time course of rotation rate for 300 sec using 970 nm bead (green: wild type live cells, blue: ΔCheY live cells). The positive and negative speed represent CW and CCW rotation, respectively. *Inset*: *y*-*x* plot of a bead rotation. Grids represent 500 nm. *Right*: Histogram of rotation rate of each graph. (b) Histograms of CW bias. The peak and SD are 0.29 ± 0.11 in wild type live cells (n = 46).

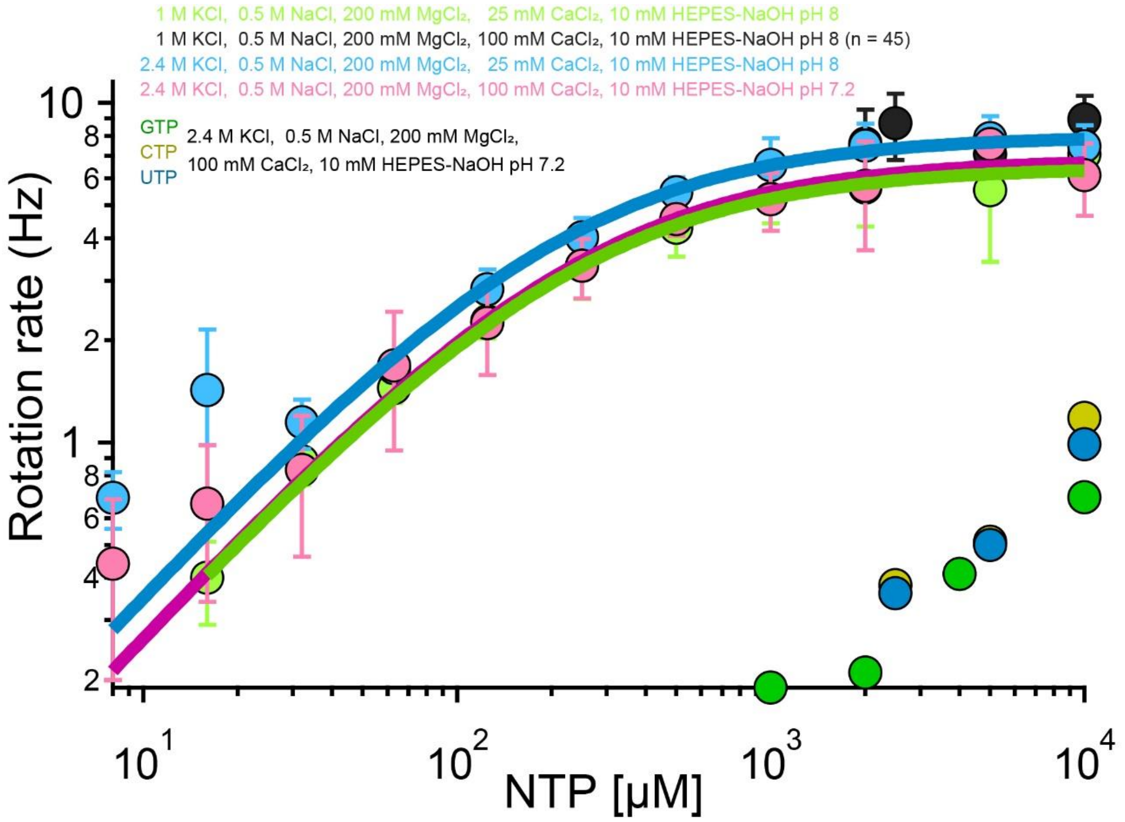

**Supplementary Figure 11 Michaelis-Menten kinetics**

The rotation rate of 500 nm bead attached to an archaellar filament. The buffer condition is indicated. The solid lines show a fit with Michaelis–Menten kinetics; $V_{\mathrm{max}}$, $K_{\mathrm{m}}$ and n are 6.47 Hz and 239 μM for light green (n = 149), 7.99 Hz and 219 μM for light blue (n = 187), and 6.83 Hz, 249 μM for red (n = 438), respectively. The data points are 62 for GTP, 63 for CTP, and 48 for UTP. Data are representative of at least two independent experiments.  We note the possibility that the observed rotation in GTP, CTP and UTP buffers might be due to contaminating ATP at a level of 1 part in ~1000, rather than rotation driven by NTPs other than ATP.

### Supplementary Tables

Table 1. Strains and plasmids

| Strain/plasmid | Relevant Phenotype/ Genotype | Note |
|---|---|---|
| ***Strains*** | | |
| ***Haloferax* (Hfx)*. volcanii* H26** | Wild type (*ΔpyrE2*) | (1) |
| ***Hfx. volcanii* FYK1** | *ΔpilB3ΔpyrE2* | This study |
| ***Hfx. volcanii* FYK2** | *ΔpilB3ΔflgA1ΔpyrE2* | This study |
| ***Hfx. volcanii* FYK3** | *flgA1(A124C)::ΔpilB3ΔflgA1* | This study |
| ***Hfx. volcanii* FYK4** | *ΔpilB3ΔflgA1ΔcheYΔpyrE2* | This study |
| ***Hfx. volcanii* FYK5** | *flgA1(A124C)::ΔpilB3ΔflgA1ΔcheY* | This study |
| ***Plasmids*** | | |
| **pTA131** | pyrE2 marked deletion plasmid. Amp resistance | (1) |
| **pSVA12703** | KO plasmid for PilB3 | This study |
| **pSVA12704** | KO plasmid for CheY | This study |
| **pSVA12727** | KO plasmid for FlgA1 | This study |
| **pSVA1392** | Tryptophan-inducible gene expression of protein | (2) |
| **pSVA5622** | Expression plasmid of FlgA1 | This study |
| **pSVA5642** | Expression plasmid of FlgA1(A124C) | This study |

Table 2. Primers

| Name | 5’ > 3’ | Note |
|---|---|---|
| **10019** | GCGGTACCCGGTGATACTGTCGATGTAGACG | Forward primer for up stream of $pilB_3$ in KO plasmid |
| **10020** | GCGGATCCGGTCGCACAGTACCTGTATCTG | Reverse primer for up stream of $pilB_3$ in KO plasmid |
| **10021** | GCGGATCCTCCCCTCGTCTCGCTCGTCGTATCTC | Forward primer for down stream of $pilB_3$ in KO plasmid |
| **10022** | GCTCTAGACGTCAATCTCGACGAGAGCTTCGC | Reverse primer for down stream of $pilB_3$ in KO plasmid |
| **10023** | GCGGTACCGTCGTCGTCCGTTCGAATAGTC | Forward primer for up stream of *cheY* in KO plasmid |
| **10024** | GCGGATCCATGAATCTCGACGTCCGGTC | Reverse primer for up stream of *cheY* in KO plasmid |
| **10025** | GCGGATCCACCTCCAGATTTCCGAAAATCACACC | Forward primer for down stream of *cheY* in KO plasmid |
| **10026** | GCTCTAGAACTTCACGGCGGTAGACGTTCG | Reverse primer for down stream of *cheY* in KO plasmid |
| **10080** | TATAGGTACCGAGCGCGGTGGTGAC | Forward primer for up stream of *flgA1* in KO plasmid |
| **10081** | GCGCGGATCCAGATTTCGTGGGTTTGGTC | Reverse primer for up stream of *flgA1* in KO plasmid |
| **10082** | GCGCGGATCCGGAGATTCAAATGTTCAACAACATC | Forward primer for down stream of *flgA1* in KO plasmid |

| | | |
|---|---|---|
| **10083** | GCTATCTAGACTGGGTCGCGAAAATGCGCAAGC | Reverse primer for down stream of *flgA1* in KO plasmid |
| **8899** | GGAATTCCATATGTTCGAAAACATCAACGAAGACC | Forward primer for ampkication of *flgA1* from *Hfx. volcanii* gDNA |
| **9054** | CGCGGATCCTCAGAGCGCAATGGGGTC | Reverse primer for ampkication of *flgA1* from *Hfx. volcanii* gDNA |
| **9098** | CGACACGTGCGACCCGGCTAACCTG | Forward primer to insert the cysteine mutation into *flgA1*(A124C) |
| **9099** | CCGGGTCGCACGTGTCGGAGCC | Reverse primer to insert the cysteine mutation into *flgA1*(A124C) |

Table 3. Population and CW bias

| **Type (number)** | **Record time** | **CW population'** | **CCW population** | **Switching population** | **CW bias** |
|---|---|---|---|---|---|
| WT live cell (47) | 5 min | 0 % (0) | 15 % (7) | 85 % (40) | 0.29 |
| ΔCheY live cell (54) | 5 min | 4 % (2) | 85 % (46) | 11 % (6) | 0.07 |
| WT ghost (61) | 10 sec | 38 % (23) | 62 % (38) | 0 % (0) | 0.35 |
| WT ghost (29) | 5 min | 24 % (7) | 66 % (19) | 10 % (3) | 0.27 |

#### Captions for Supplementary Videos

Supplementary Video 1
Swimming motility of *Haloferax volcanii* cells observed under a phase-contrast microscope. Scale bar, 50 μm.

Supplementary Video 2
Swimming ghosts in the presence of 2.5 mM [ATP]. The density of cells was frequently decreased, indicating the permeabilization of cells. After permeabilization, the ghost still exhibited the swimming motility. Scale bar, 40 μm.

Supplementary Video 3
Effect of streptavidin on a bead rotation of a live cell. At 8 sec, the buffer containing 0.1 mg $ml^{-1}$ streptavidin was added. Scale bar, 5 μm.

Supplementary Video 4
Preparation of rotary ghosts. Live cell exhibited the rotation of 500-nm bead attached to an archaellar filament. At 7 sec, the buffer was replaced with a buffer containing 0.03 % sodium cholate hydrate and 1 mg $ml^{-1}$ DNase. The density of cells was gradually decreased, indicating the permeabilization of cells. At 37 sec, the cell showing a bead rotation was permeabilized, and its motor rotation was finally stopped. At 38 sec, the buffer was then replaced with buffer containing 500 μM [ATP]. At 44 sec, the motor reactivated and exhibited rotation again. Scale bar, 20 μm.

Supplementary Video 5
ATP-dependent rotation of an archaeal motor. Bead attached to an archaellar filament rotates in the presence of 2 mM [ATP]. At 7 sec, the buffer was replaced with buffer containing 16 μM [ATP], and motor rotation was slow. At 39 sec, the buffer was again replaced with buffer containing 2mM [ATP], and the fast rotation could be seen. At 63 sec, no ATP solution was added into a chamber, demonstrating no rotation. At 90 sec, 500 μM [ATP] was added into the chamber, and the motor exhibited rotation again. Scale bar, 3 μm.

Supplementary Video6
1000-nm bead rotation of ghost at 2.5 mM [ATP]. Scale bar, 2 μm.
